# Towards high spatially resolved microproteomics using expansion microscopy

**DOI:** 10.1101/2021.03.03.433765

**Authors:** Lauranne Drelich, Soulaimane Aboulouard, Julien Franck, Michel Salzet, Isabelle Fournier, Maxence Wisztorski

## Abstract

Expansion microscopy is an emerging approach for morphological examination of biological specimens at nanoscale resolution using conventional optical microscopy. To achieve physical separation of cell structures, tissues are embedded in a swellable polymer and expanded several folds in an isotropic manner. This work shows the development and optimization of physical tissue expansion as a new method for spatially resolved large scale proteomics. Herein, we established a novel method to enlarge the tissue section to be compatible with manual microdissection on regions of interest and to carry out MS-based proteomic analysis. A major issue in the Expansion microscopy is the loss of proteins information during the mechanical homogenization phase due to the use of Proteinase K. For isotropic expansion, different homogenization agents are investigated, both to maximize protein identification and to minimize protein diffusion. Better results are obtained with SDS. From a tissue section enlarge more than 3-fold, we have been able to manually cut out regions of 1mm in size, equivalent to 300µm in their real size. We identified up to 655 proteins from a region corresponding to an average of 940 cells. This approach can be performed easily without any expensive sampling instrument. We demonstrated the compatibility of sample preparation for expansion microscopy and proteomic study in a spatial context.

**Abstract graphic:** 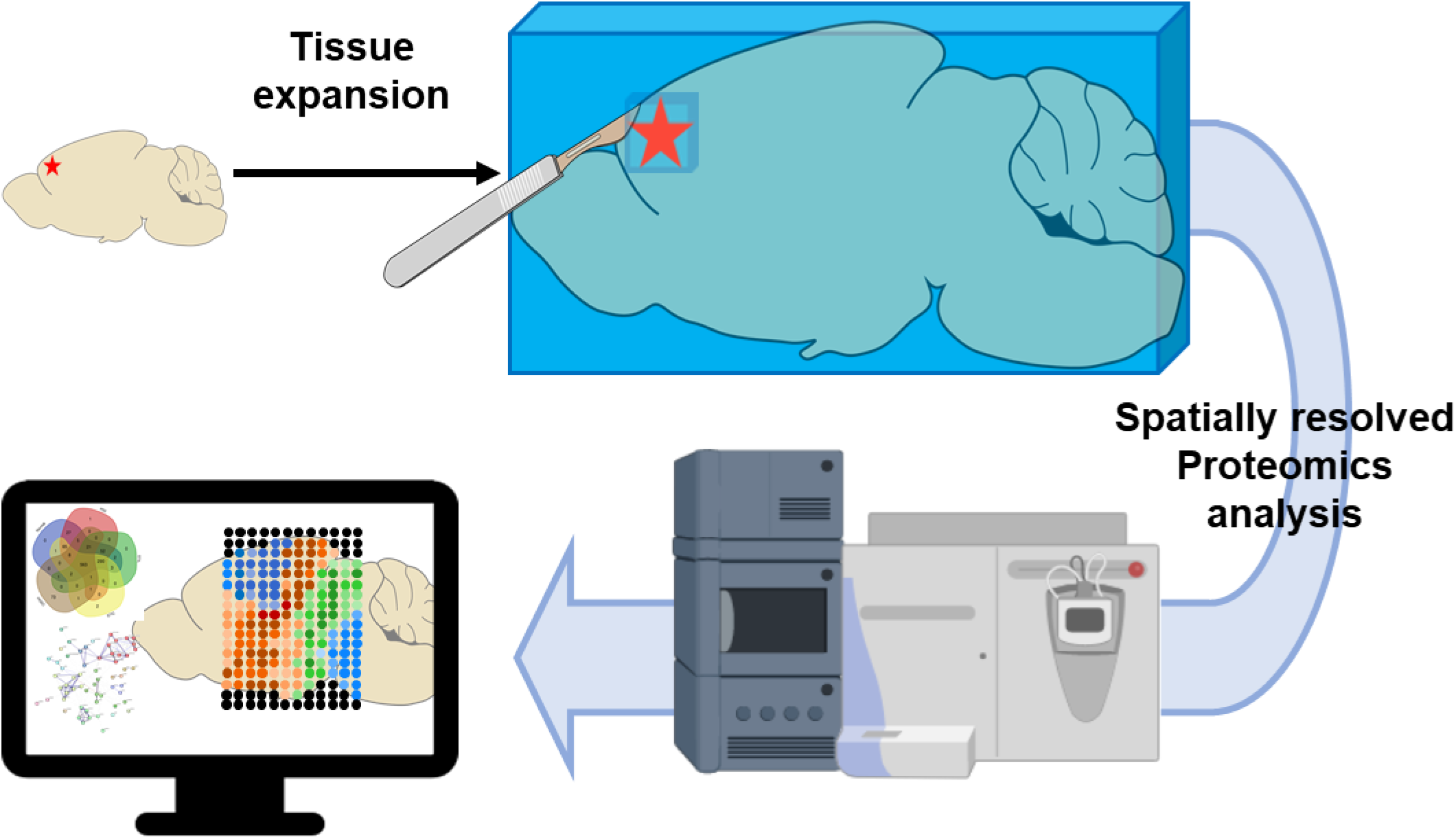

## Introduction

Today, the study of tissue and tumor heterogeneity is an emergent issue in clinical oncology. The spatial context then becomes the principal key determinant of the cellular identity in the tissue^1^. There is a real need to precisely map cells/tissue, in order to put information in a spatial context. This field is essentially approached by the way of genomics or transcriptomics analysis^1–3^. Performing proteomics analysis remains a challenge to this type of spatial approach due to the limited tools available. In fact, traditionally, organs or tissues analysis relies on a chemical homogenization to obtain cellular population that could be analyzed. These procedures would inevitably lead to a loss of information concerning the spatial localization making impossible the understanding of the architecture of the tissue.

Mass spectrometry imaging (MSI) is a powerful tool to visualize cellular heterogeneity on a tissue and could be used to determine a distinction between samples different ^4,5^. Information obtained served to guide spatially resolved analysis to identify molecules contains on these highlighted regions. In this context, several micro-sampling approaches were developed including e.g. Laser Capture Microdissection (LCM) ^6,7^, Liquid MicroJunction (LMJ) micro-extraction ^8–10^or hydrogel discs containing trypsin^11,12^. Still, their spatial accuracy is inevitably limited by the sample size that can be obtained, typically hundreds of micrometers.

Another way is to physically modify the tissue section to attain smallest regions. It has been employed for example to increase the spatial resolution that could be obtain in MALDI-MSI by stretching the sample. In this method samples are prepared by adhering tissue section to glass beads array fixed to a stretchable membrane^13^. When the membrane is stretched, it separates the tissue section into thousands of cell-sized pieces of tissue, then analyzed by MALDI-MSI.

To assess the spatial variability, we present in this paper, the development of a new way to perform spatially resolved proteomics. The originality of our approach will be to expand a piece of tissue or a tissue section in 3 dimensions to easily select regions of interest to analyze. This will be performed without any sophisticated or cost expansive instrument. We adapt the protocol developed by Boyden et al. of expansion microscopy ^14–16^in which a tissue section is magnified by 3-to 100-fold. This technique was developed initially to observe sample at a nanoscale precision with conventional microscopes. To obtain this expansion, a dense and uniform mesh of swellable polymer is introduced in the tissue. Proteins are anchored to the polymer network and the resulting hydrogel-tissue hybrid will expanse after immersion in water, resulting in physical magnification of the tissue. After some adjustments, we demonstrated that is possible to cut off a specific expanded region and treat the sample to perform MS-based proteomics analysis. Herein, we report a protocol to achieve a 3-fold volumetric expansion and obtain the identification of over 650 proteins for a region of 460µm original diameter. By doing so, we have overcome some limitation of the existing spatially resolved proteomics methods and we will be able to focus our analysis on small regions of interest on tissue section and with a limited quantity of material/cells.

## Material &Methods

The complete protocols are detailed in supporting information

### Reagents and Chemicals

For the different experiments, high purity chemicals were purchased from various supplier (complete list in the supporting information) and used without further modification.

### Tissue expansion

#### Tissue preparation

Adult Wistar male rats (University of Lille) were sacrificed according to animal use protocols approved by the university of Lille Animal Ethics committee. FFPE and Fresh frozen brain sections of 12µm thickness were used.

#### Tissue gelation

Tissue expansion was realized in accordance with the protocol published by Tillberg^14^. Complete workflow is represented in **Figure 1A**. Briefly, succinimidyl ester of 6-((acryloyl)amino)hexanoic acid (acryloyl-X (AcX), 0.1mg/mL in PBS) is deposited on tissue and incubated overnight in a humid chamber at room temperature. Gelling solution is freshly prepared. The final concentrations of chemicals in PBS (1×) are 0.01% (w/v) 4-hydroxy-2,2,6,6-tetramethylpiperidin-1-oxyl (4-Hydroxy TEMPO), 2M NaCl, 8.6% (w/v) sodium acrylate, 30%(v/v) acrylamide/bis-acrylamide (30% solution; 37.5: 1), 0.2% (w/v) TEMED, and 0.2% (w/v) APS. This solution is deposited on the tissue and spread inside a gelation chamber with a glass cover and placed in a humid chamber a 37°C for 2hours.

**Figure 1.**
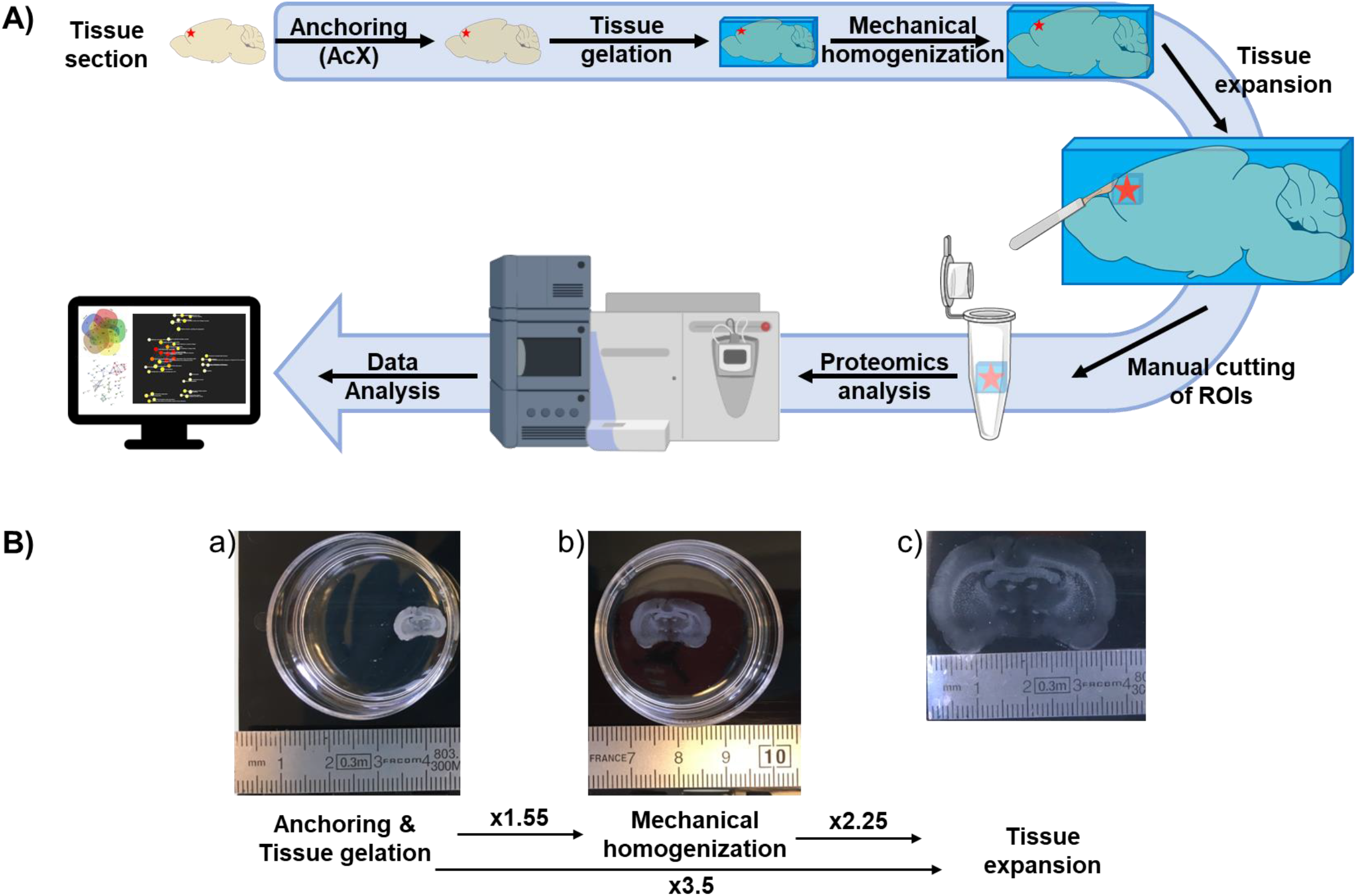
General workflow of the micro-proteomic protocol based on tissue expansion A). After tissue expansion, micro-proteomic analysis is performed by cutting a region directly in the gel using a scalpel blade. Gel-embedded tissue pieces are then subjected to enzymatic digestion and peptides are extracted to be analyzed by mass spectrometry. B). Rat brain tissue section in its original size of 1,2cm, before expansion and B. post-expansion with an expansion factor around 3 (post-expansion size of 3,8cm).

#### Mechanical homogenization

Original protocol used proteinase K (8 units/mL) diluted in a homogenization buffer (50 mM Tris/HCl (pH8), 1 mM EDTA, 0.5% Triton X-100, 1 M NaCl). Gel containing tissue is submerged in 2mL of proteinase K for 3 hours at 60°C. Several homogenization processes are tested to replace proteinase K. Different enzymes are used and to minimize the total amount of enzyme, the homogenization solution is deposited directly on the tissue section instead of a complete immersion. Our first experiment was done using a proteinase K solution diluted to 4 units/mL. In a second time, we tested several other enzymes or homogenization agent. LysC (20µg/mL in 6M Urea, 1mM EDTA) is deposited with a sufficient volume to cover the surface of the tissue section (typically 500 µl) and placed at 37°C overnight. As a second enzyme, 1mL of Trypsin-EDTA solution used for cell culture dissociation (0.025% trypsin and 0.01% EDTA in PBS) is deposited and placed at 37°C for 20 minutes. Finally, we also tested SDS as an homogenization agent, realized in first instance by adding 5% SDS diluted in the homogenization buffer (50mM TrisHCl (pH8), 1mM EDTA, 0,5% Triton X-100, 1M NaCl) and placed 30 minutes at 95°C and in a second test by decreasing the temperature at 58°C in a humidity chamber overnight.

#### Expansion and cutting

Tissue sections embedded in the gel were separated from the glass slide by submersion in HPLC water for one hour. The water is replaced 3 times every 15 minutes. Once maximal expansion size has been reached, square areas of 1×1mm^2^ or 5×5mm^2^ are cut manually with a scalpel blade and transferred into a microcentrifuge tube (LoBind 1.5mL microcentrifuge tubes, Eppendorf). A control region, located outside the tissue, is also cut to evaluate possible diffusion of proteins and peptides in the gel. The expansion factor, determined from the pre-expansion measurement (**Figure 1Ba**) divided by the post-expansion measurement (**Figure 1Bc**), is usually between 2.9- and 3.4-fold. Thus, an area of 5×5mm^2^ post-expansion corresponds to 1.6×1.6mm^2^ as a real size and around 300×300μm^2^ for a post-expanded region of 1×1mm^2^. Thereafter, results will be expressed considering the region size post-expansion: 5×5mm^2^ and 1×1mm^2^. Biopsy punch with a diameter of 1.5mm were tested for sampling regions after tissue expansion. For imaging-like analysis, 15 consecutive regions of 1×1mm^2^ were cut, each along a line through the tissue in order to assimilate each piece of gel to an image pixel.

### Proteomics analysis

Pieces of gel in the microcentrifuge tube are covered with 50μL of NH_4_HCO_3_ (50mM). Reduction process is achieved by incubation of the pieces by adding DTT solution (45mM in NH_4_HCO_3_ 50mM) 15min at 50°C. For alkylation process, gel pieces are incubated with IAA (100mM in NH_4_HCO_3_ 50mM) 15min at room temperature in obscurity. Finally, 20μL of trypsin solution (20μg/mL in NH_4_HCO_3_ 50mM) is added in each sample and incubated overnight at 37°C. Digestion is stopped by adding TFA 1% final volume.

### Liquid Microjunction experiment

To compare to another spatially resolved proteomic method, experiments of droplet-based liquid microextraction were performed according the protocol previously published^17,18^. Briefly, region of interest was first digested using trypsin solution deposited by a piezoelectric microspotter Chemical Inkjet Printer (CHIP-1000, Shimadzu, CO, Kyoto, Japan) and peptides were extracted using the TriVersa Nanomate platform (Advion Biosciences Inc., Ithaca, NY, USA) with a Liquid Extraction Surface Analysis (LESA) option.

### NanoLC-MS &MS/MS analysis

Desalting samples were analyzed by nanoLC-MS/MS (Q Exactive mass spectrometer, Thermo Scientific) (see Support Information for complete description). Proteins identification and label free quantification were performed with MaxQuant ^19,20^(Version 1.6.1). The search was done against a database containing reviewed proteome for Rattus norvegicus from UniprotKB/Swiss-Prot (8,168 sequences, July 2019). False discovery rates lower than 1% were set at peptides and proteins level. Relative label-free quantification of proteins was conducted into MaxQuant using the MaxLFQ algorithm ^21^with default parameters. The match between run (MBR) feature, with a match window of 0.7 min and an alignment window of 20 min, was activated to increase peptide/protein identification of small samples.

Analysis of identified proteins was performed using Perseus software (http://www.perseus-framework.org/) (version 1.6.0.7). The file containing the information from the identification were used and hits from reverse database, proteins with only modified peptides and potential contaminants were removed. The different methods were evaluated in terms of overlap of protein identifications (Venn diagrams) and expressed as Pearson correlation coefficients (dot plots and r value). Visualization in Venn Diagram was performed with BioVenn (http://www.biovenn.nl/) ^22^. For quantification-based mass spectrometry imaging, LFQ values of proteins were used to construct images with TIGR Multiexperiment viewer (MEV v 4.9).

The datasets used for analysis were deposited at the ProteomeXchange Consortium ^23^(http://proteomecentral.proteomexchange.org) via the PRIDE partner repository^24^with the dataset identifier PXD021919.

## Results and discussion

### Proteinase K homogenization does not allow proteins identification

On this article, we propose to develop a new methodology to perform Spatially Resolved Proteomics. To achieve this goal, we want to magnify a tissue section using a swellable polymer allowing an isotropic expansion. In first instance, we based our protocol of tissue expansion according to the publication of Tillberg ^14^concerning the protein-retention expansion microscopy (proEXM). The expansion process is divided into several phases **(Figure 1A)**. Briefly, FFPE tissue sections were deparaffinized and then cover by a solution of AcX. AcX is a linker that bind the primary amine groups on proteins and is then incorporated into the swellable hydrogel polymer during the process of gelation. The link created by AcX on the proteins will be tethered to the hydrogel polymer chains allowing the biomolecules to retain their spatial organization relative to one other. Thereafter, mechanical homogenization step consists to suppress mechanical properties in the sample in order to keep structural integrity and sample organization during expansion. Disruption of the embedded sample is generally realized by using an enzymatic digestion with the proteinase K, and the gel is finally submerged in water until maximal expansion **(Figure 1B)**.

From a post-expansion FFPE tissue section, we proceeded with a scalpel blade to a manual cutting of small squares of 5×5 mm^2^ size on the embedded tissue and outside the tissue (hydrogel alone) as a control to assess possible molecule diffusion within the gel. The gel pieces were then submitted to a conventional proteomics digestion protocol and retrieved peptides were analyzed by mass spectrometry. On the tissue, 14 proteins were identified and 9 proteins in the control region **(Figure S1)**. The number of proteins identified is low compared to what is expected for this tissue surface and an equivalent number of proteins is identified outside the tissue section. The detection of peptides outside the tissue could indicate a loss of some peptides produced during the homogenization step using the proteinase K. In fact, as FFPE tissue was used, proteins are normally crosslinked by the paraformaldehyde, preventing any diffusion within the gel. The AcX will also create some link between proteins and the hydrogel. Indeed, the proteinase K preferentially cleaves at aliphatic of aromatic amino acid residues with low specificity and leads to the formation of small peptides non-anchored to the gel. These small peptides can diffuse through the reticulation of the hydrogel during the different processing steps of expansion or proteomics analysis. This diffusion could lead to a loss of proteins localization and significantly reduce protein identification.

Nevertheless, some proteins could be identified from expanded tissue. These results incite us to investigate the replacement of the proteinase k during the mechanical homogenization step to reduce possible loss of proteins.

### Comparative analysis of different alternatives for Proteinase K

As other homogenization strategies, we tested two enzymes with specific cleavage sites, *i.e.* the trypsin and the LysC, and an anionic detergent, the SDS, which allows to disrupt protein bonds inducing protein linearization. Using the trypsin or LysC, the expansion factor was estimated to 2.1-fold; 2.3-fold with SDS and 2.4-fold with proteinase K **(Figure 2A, B, C and D)**. The correlation between the size of the sampled regions and their real size according to the expansion factor is presented in the **Table 1** with an estimation of the corresponding number of cells. For trypsin and LysC, an incomplete homogenization is observed with a lot of cracks and tissue distortion appearing during the expansion process **(Figure 2B and C)**. Using SDS, the expansion is more homogeneous **(Figure 2D)**, and the expansion factor is quite similar than the one obtained with the proteinase K.

**Table 1.**
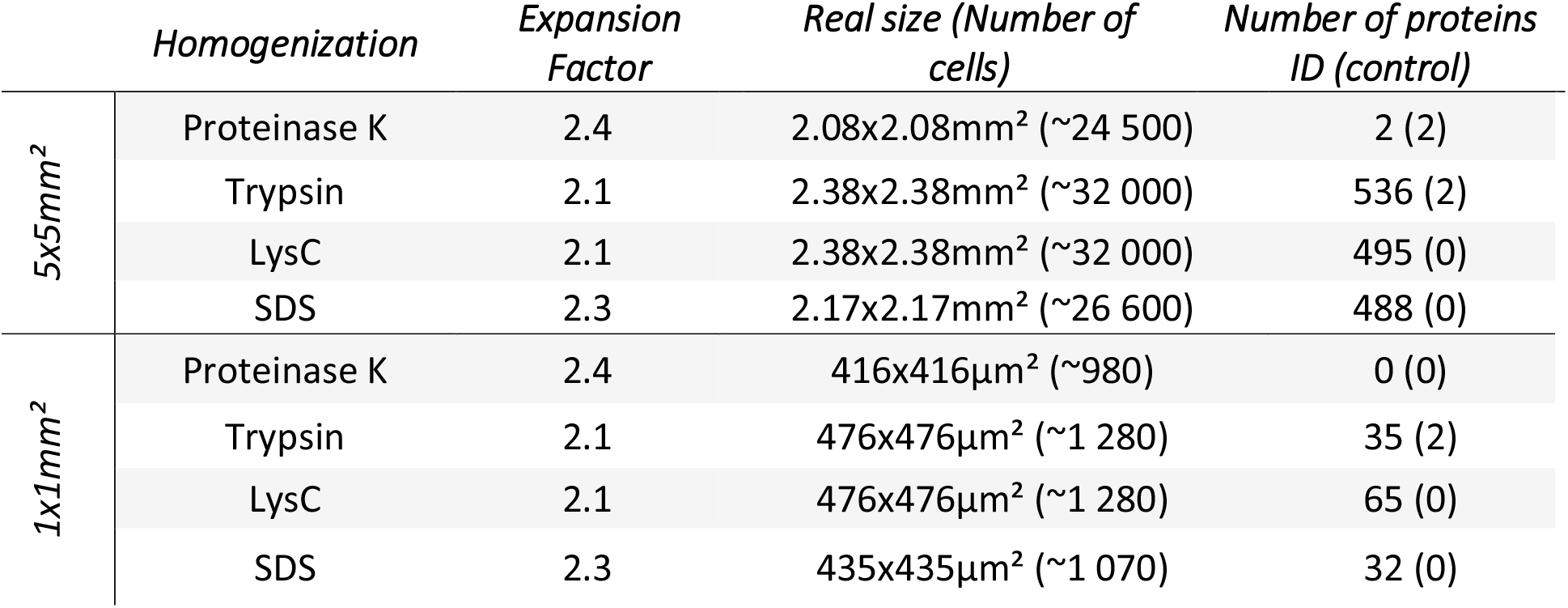
Expansion factor and number of proteins identified according to the homogenizing agent.

|  | *Homogenization* | *Expansion Factor* | *Real size (Number of cells)* | *Number of proteins ID (control)* |
| --- | --- | --- | --- | --- |
| *5x5mm<sup>2</sup>* | Proteinase K | 2.4 | 2.08x2.08mm<sup>2</sup> (~24 500) | 2 (2) |
|  | Trypsin | 2.1 | 2.38x2.38mm<sup>2</sup> (~32 000) | 536 (2) |
|  | LysC | 2.1 | 2.38x2.38mm<sup>2</sup> (~32 000) | 495 (0) |
|  | SDS | 2.3 | 2.17x2.17mm<sup>2</sup> (~26 600) | 488 (0) |
| *1x1mm<sup>2</sup>* | Proteinase K | 2.4 | 416x416µm<sup>2</sup> (~980) | 0 (0) |
|  | Trypsin | 2.1 | 476x476µm<sup>2</sup> (~1 280) | 35 (2) |
|  | LysC | 2.1 | 476x476µm<sup>2</sup> (~1 280) | 65 (0) |
|  | SDS | 2.3 | 435x435µm<sup>2</sup> (~1 070) | 32 (0) |

**Figure 2.**
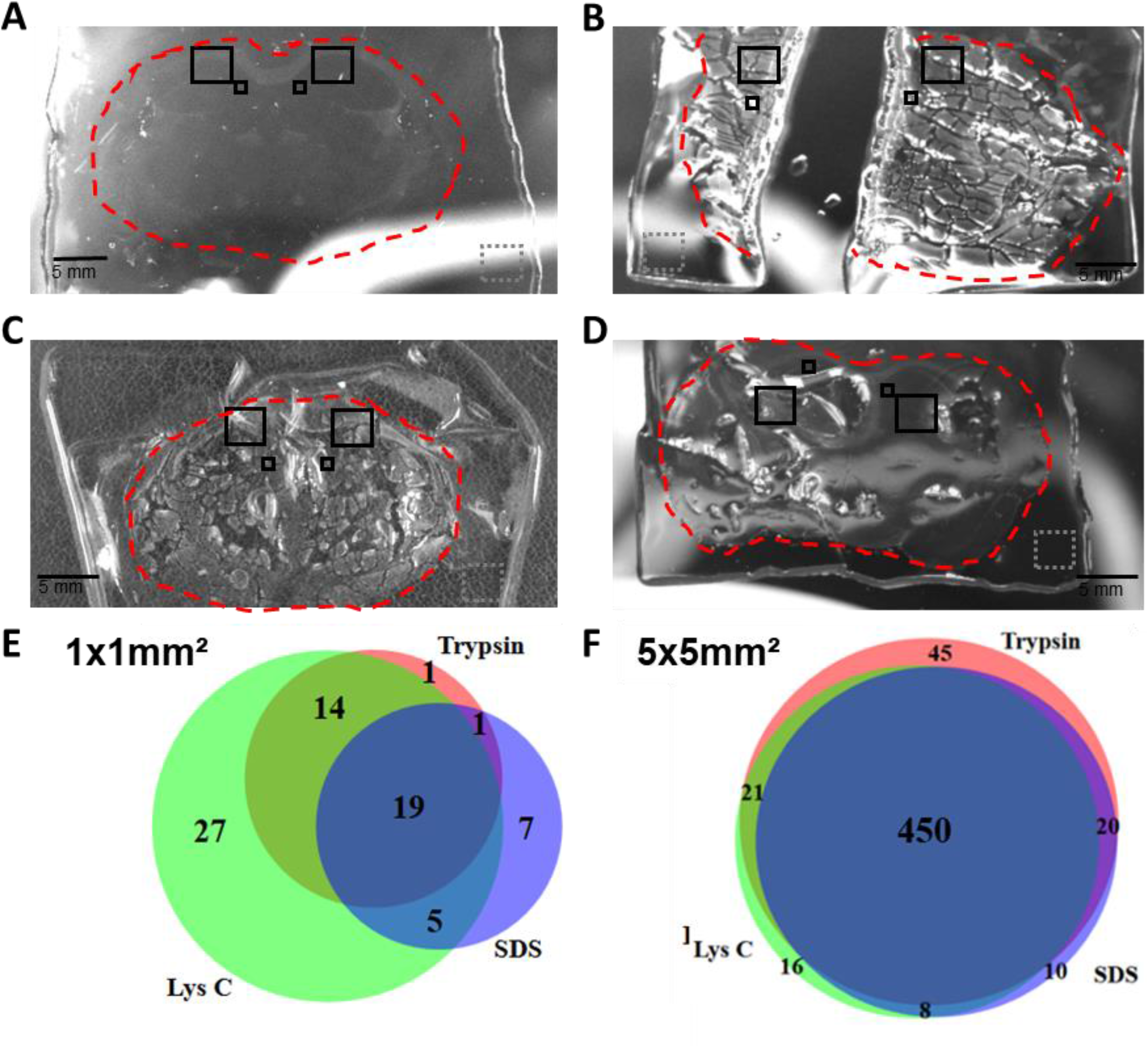
Comparative analysis of different alternatives for Proteinase K. Post-expansion rat brain sections with different homogenization processes using A) Proteinase K, B). Trypsin, C) LysC and D) SDS. The red dotted line delimits the contour of the tissue section. Squares correspond to the different sampled regions: 5×5mm^2^ and 1×1mm^2^ inside the tissue and a control area (5×5mm^2^) in the surrounding gel only. E) Venn diagrams representation of number of proteins identified using the different homogenizing agent for E) 5×5 mm^2^ square and F) 1×1 mm^2^.

For proteomics analysis, same results as above are obtained using Proteinase K with a low number of proteins identified **(Table 1)**For the other conditions, the number of identified proteins increased significantly **(Table 1 and DataS1)**. In the 5×5mm^2^ regions, up to 536 proteins were quantified using trypsin as homogenizing agent, 488 for SDS and 495 for Lys C. For 1×1mm^2^ regions, 35 proteins were quantified for trypsin, 32 proteins for SDS and 65 protein with Lys-C homogenization. In control, only 2 proteins were quantified for LysC, 1 for trypsin and none for SDS, revealing no or only very minimal diffusion within the gel.

However, as homogenization with trypsin and LysC was not homogeneous, it is difficult to precisely estimate the real size of the digested region **(Figure 2B and C)**. Based on the measured expansion factor, there is a variation of about 20% in the number of cells analyzed between the different tests **(Table 1)**. This number of cells is also difficult to estimate when the expansion is not isotropic as for trypsin and LysC. In these cases, a high number of cracks and apparent deformations were observed, which means that some groups of cells retain their original size, resulting in a higher than expected cell density in the sampled region **(Figure 2B and C; Table 1)**. This variability was also observed by comparing the proteins identified in each condition for the region of 1×1mm^2^ **(Figure 2E)**. However, for the region of 5×5 mm^2^, a high protein identification overlap (approximately 73%) was observed between Trypsin, Lys C and SDS **(Figure 2F).**

Considering these results, we decided to focus our development on SDS to replace proteinase K in the homogenization process. As a homogenized agent, SDS allows lipid removal, keep protein integrity and affect only their conformation and charge without disturbing their anchoring to the hydrogel ^14,25^.

### Method optimization and reproducibility for proteomics analysis using the SDS homogenization protocol

As a first step, we increased the homogenization time up to one night in a humid chamber with a temperature maintained to 58°C instead of 95°C to avoid liquid evaporation and hydrogel degradation. These improvements result in a greater expansion factor up to 3 times the initial tissue size.

Reproducibility was assessed by comparing triplicate sampled regions on consecutive FFPE tissue sections (S1, S2 and S3) of the midbrain **(Figure 3A)**. Only 1 protein was identified in control area from S3, no protein was identified in S1 and S2 control areas which means that proteins keep their location within the tissue despite the longer preparation time **(DataS2)**. Overlapping protein identification (Venn diagrams) was used to evaluate qualitative reproducibility and Pearson’s correlation coefficients (dot plots and r value) for quantitative reproducibility. The number of proteins identified in all replicates, regardless of their abundance, is presented in the Venn diagrams, while the quantitative values (LFQ values) were used to calculate the scatter plots.

**Figure 3.**
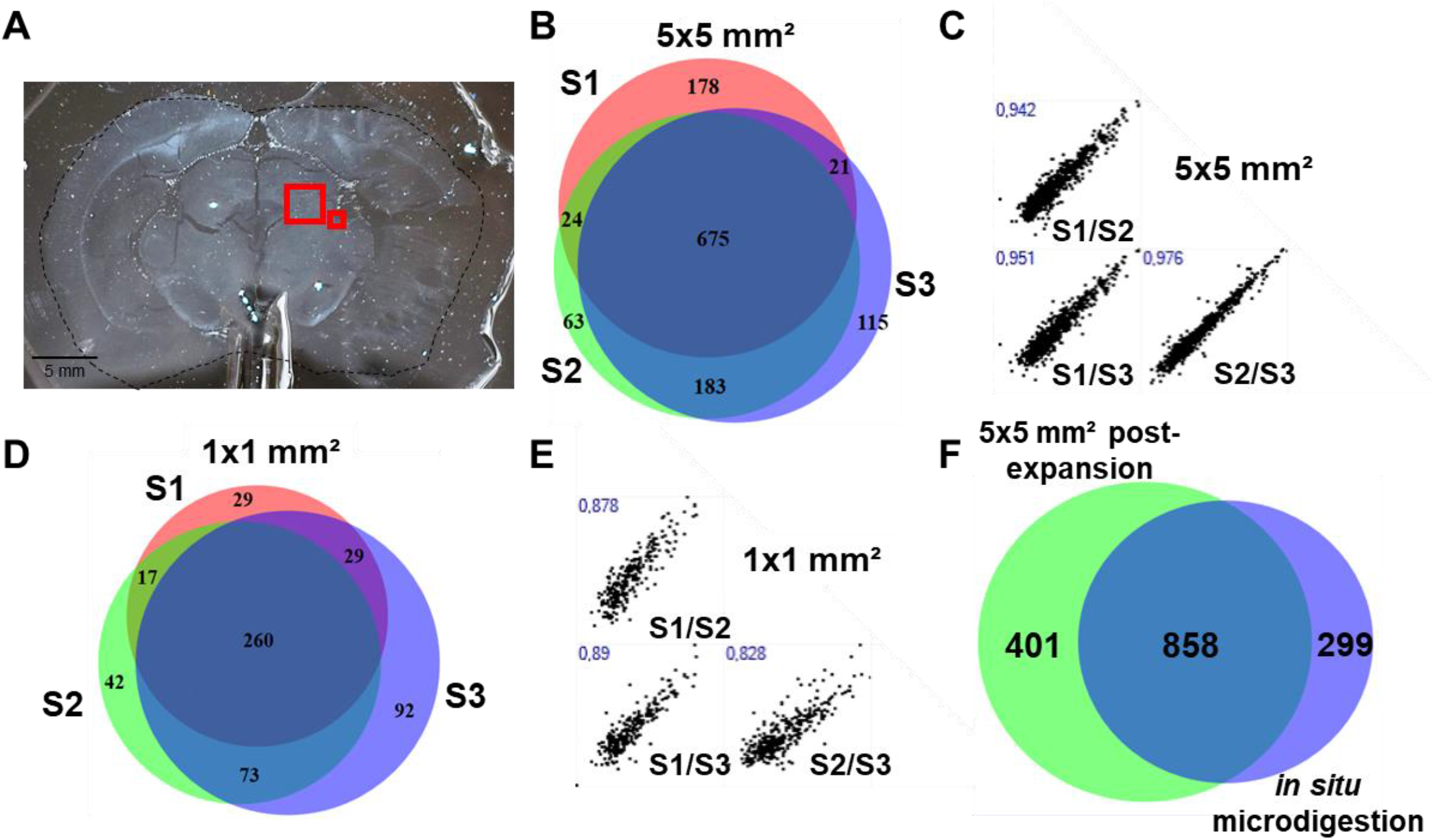
Analysis of proteomic reproducibility using the SDS homogenization protocol. A) Post-expansion widefield image of rat brain section after SDS homogenization, red square stand for the analyzed region. Experiment were performed in triplicates on 3 consecutives tissue sections. Venn diagram representation of the number of proteins identified in extraction of B) 5×5mm^2^ in size and D) 1×1mm^2^. Pearson correlation of proteins quantified in extraction of (C) 5×5mm^2^ in size and (E) 1×1mm^2^. F). Comparison of proteins identified in expansion and with digestion in situ and liquid micro junction extraction.

For 5×5mm^2^ regions, 1259 different proteins are identified with 898, 945 and 994 proteins respectively for replicate 1; 2 and 3 **(Table 2; DataS2)**. Considering the expansion factor, the real analyzed area is about 1.6×1.6 mm^2^. If we assume that cells present an average size of 15 μm and are round-shaped, we can estimate at less than 15 000 the number of cells. The optimization allows doubling the number of protein identifications compared to previous experiments. In **Figure 3B**, the Venn diagram shows that 675 proteins are shared by all replicates corresponding to 53.6% of common proteins with a small individual variation (about 5%). Looking at the direct side-by-side comparison **(Figure S2A)**, replicates S2 and S3 are closed with a high overlap in protein identification (approximately 80%). The protein content of S1 is slightly different from that of the two other replicates (around 60% of overlapping). But for the quantification, the Pearson’s correlation coefficient values show that the identified proteins gave the same quantification value between the replicates (mean of 0.956) **(Figure 3C)**.

**Table 2.**
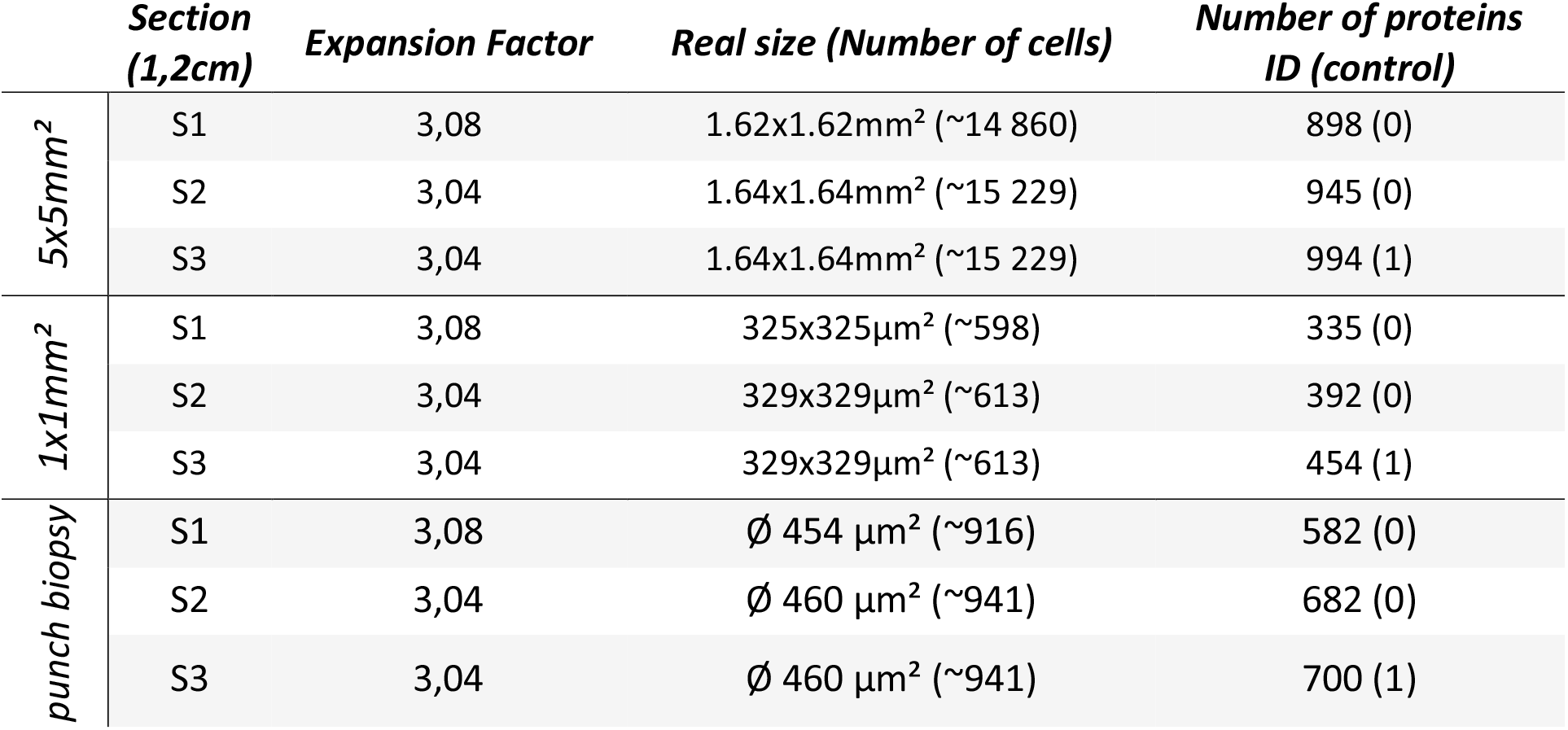
Expansion factor, estimation of number of cells and number of proteins identified after homogenization with SDS.

Reduction of the sampling region at 1×1mm^2^ after expansion correspond to a real analyzed region of 330×330 μm^2^. This surface corresponds to approximatively 600 cells and 335 groups of proteins were identified in section 1; 392 in section 2 and 454 in section 3 for a total of 542 different proteins **(Table 2; DataS2)**. A higher variability than for the 5×5mm^2^ region, was observed with 260 protein shared by all replicates representing 47.97% of common protein **(Figure 3D)** (around 60% for pairwise comparison **(Figure S2B))**. Same observations could be done for the quantitative reproducibility (variation of Pearson’s correlation coefficient from 0.83 to 0.89 **(Figure 3E)**. We observed an average of 50% of the proteins identified by the MBR feature with 5×5mm^2^ samples. It should be noted that more than 98% of the proteins are also identified in the 5×5mm^2^ regions, suggesting that using a reference sample containing a larger region or more cells for the MBR, more proteins could be identified and quantified from a smaller expanded region.

Good reproducibility both qualitatively and quantitatively is observed even though variations exist. The variation could be explained by the evolution of the histological features on the consecutive tissue sections and by the difficulties of precisely sampling the same region due to the transparency of the expanded tissue. Reproducibility of smallest regions is lower than for bigger piece of tissue, probably due to manual cutting and expansion factor variation.

### Use of fresh frozen tissue section and comparison with on tissue microdigestion and microjunction using Liquid extraction surface analysis

Formalin fixation is advantageous for preservation and conservation of cellular and architectural morphologic detail in tissue sections but results in formaldehyde cross-links and many proteins are still not detectable using these approaches ^26^. For this purpose, we applied also the strategy developed to fresh frozen tissue section with encouraging results but a poor reproducibility and great variability. Indeed, we identify a total of 1 038 proteins in 5×5mm^2^ samples in all replicates with respectively 136 proteins in section 1, 78 in section 2 and 788 in section 3 **(Table 3)**. As the tissue was not fixed, the different steps, in particular the mechanical homogenization, lead to significant diffusion of proteins within the gel resulting in a loss of the actual localization (*i.e.* 10 proteins, 26 and 0 respectively in the control region of sections 1; 2 and 3 - **Table 3)**.

**Table 3.** Expansion factor and number of proteins identified in fresh frozen tissue after homogenization with SDS in each triplicate.

|  | *Section (1,5cm)* | *Expansion Factor* | *Real size (Number of cells)* | *Number of proteins ID (control)* |
| --- | --- | --- | --- | --- |
| *5x5mm<sup>2</sup>* | S1 | 3,26 | 1.53x1.53mm<sup>2</sup> (~13 248) | 136 (10) |
|  | S3 | 3,33 | 1.50x1.50mm<sup>2</sup> (~12 733) | 78 (26) |
|  | S2 | 3,2 | 1.56x1.56mm<sup>2</sup> (~13 772) | 788 (0) |

We then performed a comparison with another direct surface sampling method using in situ digestion and extraction by liquid microextraction. This micro extraction strategy was used in particular for tissue microenvironment characterization^5,8,10,18,27^. The results obtained after tissue expansion were compared to one obtained with the best setup using this method.

On-tissue enzymatic digestion combined with liquid microextraction performed on a fresh frozen tissue section allows identification of 1157 proteins in a region of 900μm in diameter. Compared to the total of proteins identified in the replica of the same region from a region of 5×5mm^2^ post-expansion FFPE tissue section, we observed 858 commons proteins **(Figure 3F)**. Equivalent number of proteins are identified in the two techniques with 299 proteins unique to *in situ* microdigestion/liquid microextraction whereas 401 are unique to the replicate of post-expansion tissue proteomics **(DataS3)**. The observed variation is probably due to the difference in the size of the region (spot of 900μm in diameter for LESA versus 1.6×1.6 mm^2^ square for expansion), the method of digestion and the formalin fixation of the tissue used for expansion, whilst microextraction was performed on fresh frozen tissue which gave better accessibility to proteins than on fixed tissue.

### Use of a biopsy punch for small regions sampling

We observed that sampling the same region in consecutive sections with a scalpel blade may result in differences in gel size. In this sense, an instrument capable of cutting a specific region with reproducible size was subsequently used. To improve reproducibility, a biopsy punch was used to cut and remove a tissue disc. This allowed to precisely cut a 1400 μm diameter gel disc (**Figure 4A**) corresponding to ~460 μm diameter region of tissue before expansion (around 940 cells) **(Table 2)**. For three consecutive regions, we obtained punch diameter of 1400±37μm. A total of 808 distinct groups of protein were obtained in the replicates (582, 682 and 700 in replicate S1, S2 and S3 respectively (**Figure 4B; DataS2**). Venn diagram shows a good reproducibility between the three replicates with 62.13% of common protein. The quantitative reproducibility is high with an average Pearson’s correlation of 0.949 and attain 0.97 between S2 and S3 (**Figure 4C)**. For individual variability between the replicates, the same observations as for manual sampling are made with higher differences between replicates S1 and S3 **(Figure S2C)**.

**Figure 4.**
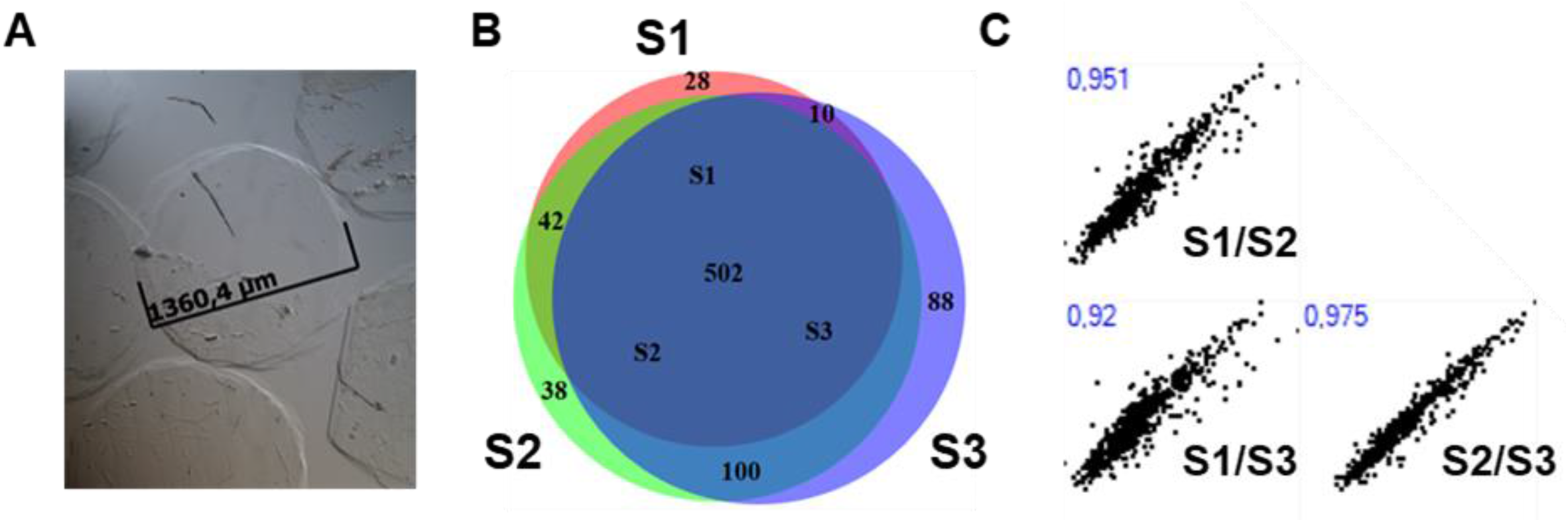
Utilization of punch biopsy improve sampling reproducibility. A) Disk of gel using a punch biopsy. B) Venn diagram representation of the number of proteins identified using punch biopsy extraction. Experiment were performed in triplicates on 3 consecutives tissue sections C) Pearson correlation of LFQ in each replicate.

These results highlight that variation observed in the proteomics content is mainly due to the manual cut of the region of interest. Indeed, it is difficult to sample gel pieces of the same size manually. Reproducibility is significantly improved with the use of a punch biopsy. This circular blade allows to cut region of same size easily and with precision. Consequently, we observed a reduction of individual variation and considerable increase of common protein and of quantitative reproducibility. A variation is still observed certainly due to minor variation in the position of the sampling due to the transparency of the tissue section. Evolution of the histological features between the tissue sections or variation of the expansion factor could also be a source of variations. Similar results were recently obtained using enzyme delivering hydrogels as a tool for localized analyte extraction directly on the tissue sample ^11,12^, allowing the identification of about 700 proteins with a hydrogel of 357μm diameter ^12^.

### Quantification-based mass spectrometry profiling using tissue expansion

Next step consisted of analyzing consecutive adjacent points to perform a quantification-based profiling, as we have done in other publications^9,28,29^. To achieve this, we sampled 15 consecutive regions of 1×1mm^2^ each along a line through a rat brain tissue section in order to assimilate each piece of gel to an image pixel **(Figure 5 A)**. It is then possible to use MS-based quantification data for an image reconstruction^28^. All extracts were digested and analyzed by LC-MS/MS. A total of 511 proteins were identified. Images were constructed and each square represents a piece of gel with its position on the tissue. The differences in expressions are color coded, red represents protein overexpression whereas dark blue represent a low detection **(Figure 5B and C, Table S1)**. Despite a spatial resolution of 300μm, expressions of proteins are different depending on the localization in the tissue. Expression of proteins specific in each region of the rat brain was then examined. The validation of this spatial mapping is confirmed by the overexpression of Creatine kinase and Calcium/calmodulin-dependent protein kinase in cortex and fimbria of the hippocampus which are known to contain these proteins ^30,31^. In these regions we also observed overexpression of Alpha-internexin relative to thalamus. This protein is an intermediate filament involved in neuron morphogenesis and neurite outgrowth ^32^. Cofilin-1 appears to be preferentially expressed in hippocampus region. This specific protein, as a regulator of actin dynamics, may contribute to degenerative processes ^33^. Preferential distribution of GFAP and Pro SAAS is observed in thalamus, which is consistent with previous publications ^28,30^. In particular, MBP quantification is in accordance with immunohistology data already published ^9^. Proteins from housekeeping genes such as Tubulin beta-4 chain and 14-3-3 protein epsilon were also identified. These proteins are known to have low variation in expression in tissues, here we observed a coefficient of variation of 2% and 1% respectively for theses 2 proteins **(Figure 5C)**.

**Figure 5.**
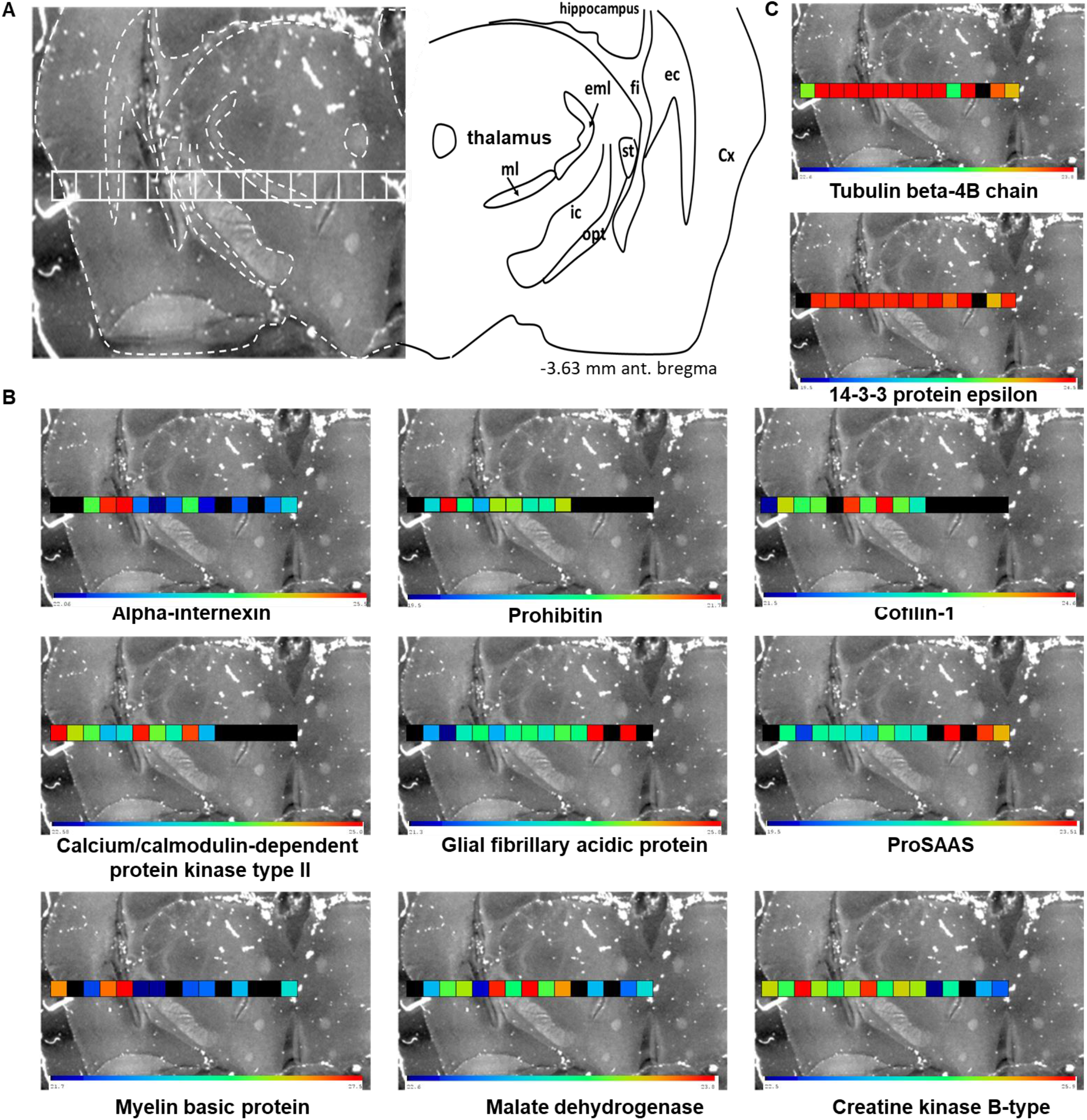
Quantification-based mass spectrometry profiling using tissue expansion A) Image of post-expansion rat brain tissue section, and histological annotations (ec: external capsule; fi: fimbria of the hippocampus; ic: internal capsule; C×: cerebral cortex; st: stria terminalis; opt: optic tract; eml: external medullary lamina; ml: medial lemniscus). Squares represent extractions for microproteomic. B) Reconstructed distribution of representative proteins and C) housekeeping genes based on label free quantification values

This strategy provides indirect molecular imaging based on the identification and subsequent quantification of proteins with a spatial resolution of less than 350 μm. It represents a new interesting feature for MSI as it enables to map many high and low abundance proteins. Recently, methods have been developed to increase the number of proteins identification from small number of cells. For example, the nanoPOTS^34^method improve the identification of thousands of proteins from a reduced number of cells. The combination of the tissue expansion with this type of methods would certainly allow an increase of the number of proteins identified from small regions.

## Conclusion

In conclusion, we demonstrated for the first time that expansion microscopy is plainly compatible with conventional large-scale mass spectrometry-based proteomics. Substituting the proteinase K used in the mechanical homogenization step by SDS reduces protein loss due to the generation of small peptides that diffuse through the hydrogel. Using our protocol, a physical expansion of FFPE tissue section by more than 3-fold was achieved, makes it possible to withdraw a well-controlled area easily and manually with an original size down to 330μm on side. We successfully used this protocol to perform spatially resolved proteomics analysis of regions of interest and to identify more than 655 proteins for an equivalent of less than 940 cells. Identification of proteins from expanded tissue showed a good qualitative and quantitative reproducibility. This strategy is particularly useful for mapping the protein content of closely related regions on a tissue section which has been challenging in previous spatially resolved proteomics studies. We performed a quantification-based mass spectrometry imaging of more than 500 proteins with a lateral resolution close to 300μm. This resolution can be further reduced by investigating some new expansion microscopy protocols like the ×10 Expansion Microscopy ^35^, ZOOM (Zoom by hydrOgel cOnversion Microscopy) ^36^or iterative expansion microscopy (iE×M) ^37^to expand biological sample up to 20-to 100-fold. Interestingly as our protocol is compatible with conventional expansion microscopy it will be possible to target a protein using fluorescent antibodies ^14^and obtain the proteome of the expression region of this protein. Tissue expansion can also be combined with ExFISH which involve the use of linker that enables RNA to be covalently attached to the gel ^38,39^. Then, fluorescent in situ hybridization (FISH) imaging of mRNA can be done ^38^and the region containing fluorescence signal can be delimited, cut and analyzed to observe the presence of the translated protein by mass spectrometry. Thus, our strategy could be used as a simple single cell tool for proteomic analysis in order to reveal variations of in limited cell population and understand specific disease mechanisms.

## Supporting information

DataS1

DataS2

DataS3

## Supporting information

Supporting information.docx: contained 2 supporting figures, 1 supporting table and supporting methods

DataS1.xlsx contained the list of quantified proteins for post-expansion region of 5×5mm^2^ or 1×1mm^2^ and off tissue using different homogenization agents

DataS2.xlsx: contained the list of quantified proteins post-expansion in reproducibility experiments using SDS as homogenization agent.

DataS3.xlsx contained the list of quantified proteins corresponding to Venn diagram for post-expansion tissue proteomics vs in situ microdigestion/liquid microjunction extraction.

## Acknowledgements

We acknowledge M. Denfer and PG. Ferron for their technical assistance. This work was funded by University of Lille, Ministère de l’Enseignement Supérieur, de la Recherche et de l’Innovation and Inserm.

