## supporting information for "Towards high spatially resolved microproteomics using expansion microscopy"

Univ. Lille, Inserm, CHU Lille, U1192 - Protéomique Réponse Inflammatoire Spectrométrie de Masse - PRISM, F-59000 Lille, France.

*

### TABLE of CONTENTS

#### Supporting files:

**DataS1.xlsx**: List of quantified proteins for post-expansion region of 5x5mm² or 1x1mm² and off tissue using different homogenization agents

**DataS2.xlsx:** List of quantified proteins post-expansion in reproducibility experiments using SDS as homogenization agent.

**DataS3.xlsx:** List of quantified proteins corresponding to Venn diagram for post-exapansion tissue proteomics vs *in situ* microdigestion/liquid microjunction extraction.

#### Supporting figures:

**Figure S1.** Protein Identification after homogenization with Proteinase K on tissue and on the surrounding gel only.

**Figure S2.** Side by side comparison between the three replicates for region of A) 5x5 mm², B) 1x1mm² and C) using a punch biopsy

#### Supporting tables:

**Table S1:** Label free quantification values (LFQ) used for the quantification-based mass spectrometry profiling using tissue expansion.

#### Supporting methods:

Reagents and chemicals
Tissue preparation
NanoLC-MS &MS/MS analysis
Data analysis

**Supporting. Figures**

**Figure S1.** Protein Identification after homogenization with Proteinase K on tissue and on the surrounding gel only.

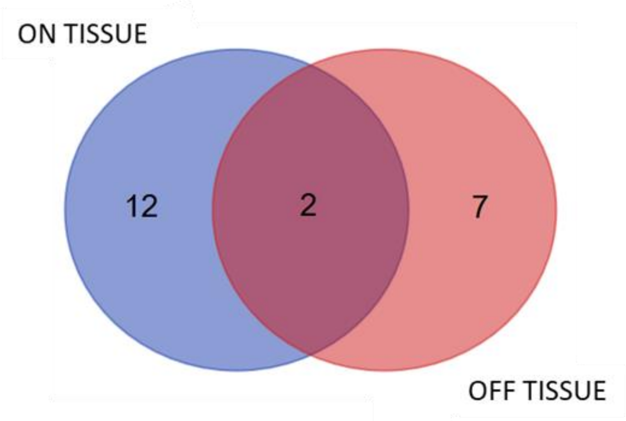


| On tissue | Off tissue |
| --- | --- |
| D3ZQG6 | P11275 |
| P02091 | P62632 |
| P07825 | P63039 |
| P0CC09 | P63259 |
| P15999 | P68370 |
| P60203 | P85108 |
| P85108 | Q5XI22 |
| Q00715 | Q5YLM1 |
| Q5XIF6 | Q6P9V9 |
| Q62634 | |
| Q6P9T8 | |
| Q6P9V9 | |
| Q8CFG6 | |
| Q9WV48 | |

**Figure S2.** Side by side comparison between the three replicates for region of A) 5x5 mm², B) 1x1mm² and C) using a punch biopsy

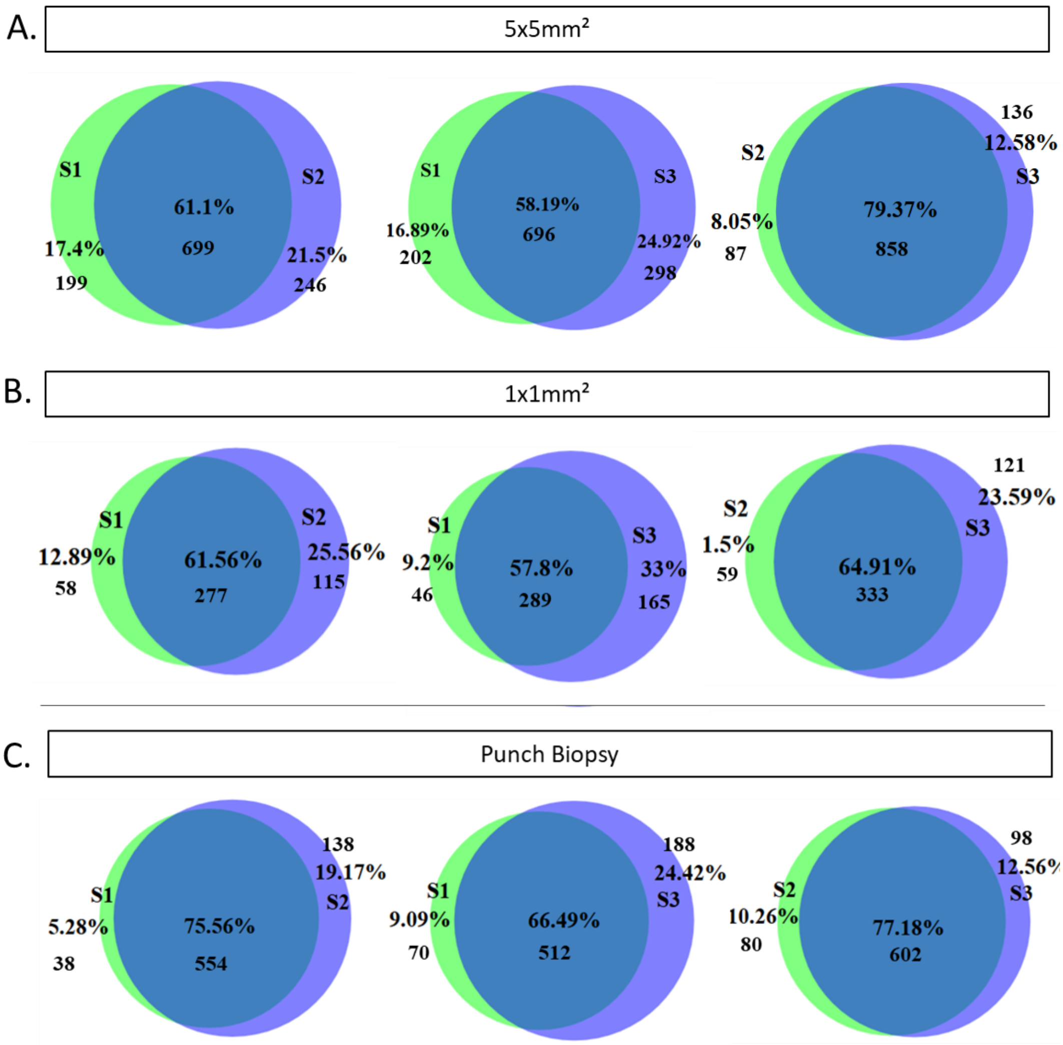

Supporting table

**Table S1:** Label free quantification values (LFQ) used for the quantification-based mass spectrometry profiling using tissue expansion.

| LFQ intensity | | | | | | | | | | | | | | | | | |
|---|---|---|---|---|---|---|---|---|---|---|---|---|---|---|---|---|---|
| LFQ intensity 1 | LFQ intensity 2 | LFQ intensity 3 | LFQ intensity 4 | LFQ intensity 5 | LFQ intensity 6 | LFQ intensity 7 | LFQ intensity 8 | LFQ intensity 9 | LFQ intensity 10 | LFQ intensity 11 | LFQ intensity 12 | LFQ intensity 13 | LFQ intensity 14 | LFQ intensity 15 | T: Majority protein IDs | T: Protein names | T: Gene names |
| 23.95 | 26.17 | 26.41 | 26.31 | 26.31 | 26.06 | 25.84 | 26.14 | 25.99 | 25.83 | 23.37 | 25.86 | NaN | 25.29 | 24.63 | Q6P9T8 | Tubulin beta-4B chain | Tubb4b |
| NaN | 24.24 | 24.09 | 24.56 | 24.3 | 24.72 | 24.22 | 24.74 | 24.16 | 24.87 | 23.95 | 24.66 | NaN | 23.47 | 24.26 | P62260 | 14-3-3 protein epsilon | Ywhae |
| 26.98 | 24.36 | 24.13 | 23.4 | 23.59 | 24.9 | 24.19 | 23.77 | 24.79 | 23.37 | NaN | 22.58 | NaN | NaN | NaN | P11275 | Calcium/calmodulin-dependent protein kinase type II subunit alpha | Camk2a |
| NaN | 22.62 | 21.33 | 23.42 | 23.92 | 22.71 | 23.75 | 23.97 | 23.37 | 24.13 | 23.84 | 25.74 | NaN | 25.7 | NaN | P47819 | Glial fibrillary acidic protein | Gfap |
| NaN | 21.72 | 19.97 | 21.6 | 21.47 | 21.24 | 20.81 | 22.02 | 21.4 | 21.31 | NaN | 23.51 | NaN | 23.23 | 22.69 | Q9QXU9 | ProSAAS;KEP;Big SAAS;Little SAAS;Big PEN-LEN;PEN;PEN-20;Little LEN;Big LEN | Pcsk1n |
| NaN | 22.06 | 24.25 | 25.31 | 25.45 | 22.78 | 22.08 | 22.81 | 24.18 | 22.4 | NaN | 22.6 | NaN | 22.84 | 23.42 | P23565 | Alpha-internexin | Ina |
| NaN | 20.38 | 21.67 | 20.98 | 20.2 | 21.07 | 21.02 | 20.56 | 20.62 | 21.11 | NaN | NaN | NaN | NaN | NaN | P67779 | Prohibitin | Phb |
| 21.56 | 23.81 | 23.39 | 23.52 | NaN | 24.38 | 23.48 | 24.54 | 23.54 | 22.94 | NaN | NaN | NaN | NaN | NaN | P45592 | Cofilin-1 | Cfl1 |
| 26.57 | NaN | 22.47 | 26.78 | 27.44 | 21.73 | 21.81 | NaN | 22.58 | 22.8 | NaN | 23.63 | NaN | NaN | 24.15 | P02688 | Myelin basic protein | Mbp |
| NaN | 22.99 | 23.37 | 23.46 | 22.69 | 23.74 | 23.3 | 23.78 | 23.36 | 23.59 | NaN | 23 | NaN | 22.83 | 23.09 | P04636 | Malate dehydrogenase, mitochondrial | Mdh2 |
| 25.04 | 24.6 | 25.84 | 24.91 | 24.53 | 24.88 | 25.68 | 24.49 | 25.07 | 24.92 | 22.52 | 24.23 | NaN | 23.55 | 23.09 | P07335 | Creatine kinase B-type | Ckb |

### Supporting Methods:

#### Reagents and chemicals

6-((acryloyl)amino) hexanoic acid (acryloyl-X or AcX), Chloroform (CHCl3), ethanol (EtOH), HPLC grade methanol (MeOH), paraformaldehyde (PFA), water and Trypsin-EDTA solution were obtained from Thermo Fisher Scientific (Courtabœuf, France). 4-hydroxy-2,2,6,6-tetramethylpiperidin-1-oxyl (4-hydroxy-TEMPO, 97%), ethylenediaminetetraacetic acid (EDTA), iodoacetamide (IAA), AR grade trifluoracetic acid (TFA, 99%), Triton X-100, sodium citrate, acrylamid/bis-acrylamid (30% ; 37.5:1) (AA/BAA), sodium acrylate (SA, 97%) and ammonium bicarbonate (NH4HCO3) were purchased Sigma-Aldrich (Saint-Quentin Fallavier, France). Xylene and formic acid (FA, ≥96%) are from Biosolve (Dieuze, France). Sequencing grade modified porcine trypsin, sequencing grade proteinase K, sequencing grade Trypsin/LysC and sequencing grade LysC were obtained from Promega (Charbonnieres, France). HPLC grade Acetonitrile (ACN) come from VWR (Fontenay-sous-Bois, France). DL-dithiothreitol (DTT) and urea were purchased from Euromedex (Souffelweyersheim, France). Sodium dodecyl sulfate (SDS), N,N,N′,N′-tetramethylethane-1,2-diamine (TEMED) and ammonium persulfate (APS) are from Bio-Rad (Marnes-la-Coquette, France).

#### Tissue preparation

Rat brain, both FFPE and fresh frozen, were cut at 12µm. A cryostat (Leica Microsystems, Nanterre, France) was used to cut the fresh frozen tissue and a microtome to cut FFPE tissue. FFPE tissue sections were mounted on SuperFrost Plus glass slide (Thermo Fischer Scientific, Courtaboeuf, France) and fresh frozen tissue on non poly-L-lysine-coated glass and stored a -80°c until use. FFPE tissues were dewaxed with xylene (2x3min), ethanol 100% (2x3min), ethanol 95% (1x3min), ethanol 70% (1x3min), ethanol 50% (1x3min) and distilled water (1x3min).

#### NanoLC-MS & MS/MS analysis

After digestion, samples were subjected to desalting using C-18 Ziptip (Millipore, Saint-Quentin-en-Yvelines, France) eluted by 80% ACN and dried under vacuum. Dried samples were reconstituted in 0.1% FA aqueous/ACN (98:2, v/v).

Two systems were used to separate peptides, a nanoACQUIITY UPLC (Waters) for samples using proteinase K for homogeneization and Imaging like strategy and a EASY-nLC 1000 (Thermo Scientific) for all other experiment. Each nanoLC was set to obtain same performance. For nanoACQUITY UPLC, samples were separated by online reversed phase using a pre-concentration column (nanoAcquity Symmetry C18, 5 µm, 180 µm x 20 mm) and an analytical column (nanoAcquity BEH C18, 1.7 µm, 75 µm x 250 mm). The peptides were separated by applying a linear gradient of acetonitrile in 0.1% formic acid (5%-35%) for 2 hours, at the flow rate of 300 nl/min. For EASY-nLC1000, samples were separated by online reversed-phase using a Proxeon trap column (75 µm ID x 2 cm, 3 µm, Thermo Scientific,) and a C18 packed-tip column (Acclaim PepMap, 75 µm ID x 50 cm, 2 µm, Thermo Scientific). The digested peptides were separated using an increasing amount of acetonitrile in 0.1% formic acid from 2 to 30% for 2 hours at a flow rate of 300 nL/min. A voltage of 2.4 kV was applied by the liquid junction in order to electrospray the eluent using the nanospray source. Each chromatography system was coupled to a high-resolution mass spectrometer Q-Exactive (Thermo Scientific). The mass spectrometer was operated in data dependent mode defined to analyze the 10 most intense ions of MS analysis (Top 10). The MS analysis was performed with an m/z mass range between 300 to 1600, a resolution of 70,000 FWHM, an AGC of 3e6 ions and a maximum injection time of 120 ms. For MS/MS analysis, the scan range was between m/z 200 to 2000, an AGC was set at 5e4 ions and the resolution was set at 17,500 FWHM. Default charge state was set at 2, unassigned and +1 charge states were rejected, HCD with a normalized energy of 30 was used, and dynamic exclusion was enabled for 25s.

**Data analysis**

All MS data were processed with MaxQuant [1,2] (Version 1.6.1 ) using Andromeda [3] search engine. Proteins were identified by searching MS and MS/MS data against reviewed proteome for Rattus norvegicus from UniprotKB/Swiss-Prot (8,168 sequences, July 2019). Trypsin specificity was used for digestion mode, with N-terminal acetylation and methionine oxidation selected as variable. Carbamidomethylation was set as a fixed modification for tissue expansion analysis. We allowed up to two missed cleavages. An initial mass accuracy of 6 ppm was selected for MS spectra, and MS/MS tolerance was set to 20 ppm for HCD data. FDR at peptide spectrum matches (PSM) and protein level was estimated using a decoy version of the previously defined databases (reverse construction) and set to 1%. Relative label-free quantification of proteins was conducted into Max-Quant using the MaxLFQ algorithm [4] with default parameters. The match between run (MBR) feature, with a match window of 0.7 min and an alignment window of 20 min, was activated to increase peptide/protein identification of small samples. Analysis of identified proteins was performed using Perseus software (http://www.perseus-framework.org/) (version 1.6.0.7).

(1) Cox, J.; Mann, M. MaxQuant Enables High Peptide Identification Rates, Individualized

p.p.b.-Range Mass Accuracies and Proteome-Wide Protein Quantification. *Nat. Biotechnol.* **2008**, *26* (12), 1367–1372. https://doi.org/10.1038/nbt.1511.

(2) Tyanova, S.; Temu, T.; Carlson, A.; Sinitcyn, P.; Mann, M.; Cox, J. Visualization of LC-MS/MS Proteomics Data in MaxQuant. *Proteomics* **2015**, *15* (8), 1453–1456. https://doi.org/10.1002/pmic.201400449.

(3) Cox, J.; Neuhauser, N.; Michalski, A.; Scheltema, R. A.; Olsen, J. V; Mann, M. Andromeda: A Peptide Search Engine Integrated into the MaxQuant Environment. *J. Proteome Res.* **2011**, *10* (4), 1794–1805. https://doi.org/10.1021/pr101065j.

(4) Cox, J.; Hein, M. Y.; Luber, C. A.; Paron, I.; Nagaraj, N.; Mann, M. Accurate Proteome-Wide Label-Free Quantification by Delayed Normalization and Maximal Peptide Ratio Extraction, Termed MaxLFQ. *Mol. Cell. Proteomics* **2014**, *13* (9), 2513–2526. https://doi.org/10.1074/mcp.M113.031591.
